# Collagen density regulates the activity of tumor-infiltrating T cells

**DOI:** 10.1101/493437

**Authors:** Dorota E Kuczek, Anne Mette H Larsen, Marco Carretta, Adrija Kalvisa, Majken S Siersbæk, Ana Micaela C Simões, Anne Roslind, Lars H Engelholm, Marco Donia, Inge Marie Svane, Per thor Straten, Lars Grøntved, Daniel H Madsen

**Author notes:** Corresponding author: Daniel Hargbøl Madsen.

## Abstract

**Background:** Tumor progression is accompanied by dramatic remodeling of the surrounding extracellular matrix leading to the formation of a tumor-specific ECM, which is often more collagen-rich and of increased stiffness. The altered ECM of the tumor supports cancer growth and metastasis, but it is unknown if this effect involves modulation of T cell activity. To investigate if a high-density tumor-specific ECM could influence the ability of T cells to kill cancer cells, we here studied how T cells respond to 3D culture in different collagen densities.

**Methods:** T cells cultured in 3D conditions surrounded by a high or low collagen density were imaged using confocal fluorescent microscopy. The effects of the different collagen densities on T cell proliferation, survival, and differentiation were examined using flow cytometry. Cancer cell proliferation in similar 3D conditions was also measured. Triple-negative breast cancer specimens were analyzed for the number of infiltrating CD8+ T cells and for the collagen density. Whole-transcriptome analyses were applied to investigate in detail the effects of collagen density on T cells. Computational analyses were used to identify transcription factors involved in the collagen density-induced gene regulation. Observed changes were confirmed by qRT-PCR analysis.

**Results:** T cell proliferation was significantly reduced in a high-density matrix compared to a low-density matrix and prolonged culture in a high-density matrix led to a higher ratio of CD4+ to CD8+ T cells. The proliferation of cancer cells was unaffected by the surrounding collagen-density. Consistently, we observed a reduction in the number of infiltrating CD8+ T-cells in mammary tumors with high collagen-density indicating that collagen-density has a role in regulating T cell abundance in human breast cancer.

Whole-transcriptome analysis of 3D-cultured T cells revealed that a high-density matrix induces downregulation of cytotoxic activity markers and upregulation of regulatory T cell markers. These transcriptional changes were predicted to involve autocrine TGF-B signaling and they were accompanied by an impaired ability of tumor-infiltrating T cells to kill autologous cancer cells.

**Conclusions:** Our study identifies a new immune modulatory mechanism, which could be essential for suppression of T cell activity in the tumor microenvironment.

## Background

Solid tumors consist of cancer cells interacting with the tumor microenvironment, which includes stromal cells, immune cells, and the extracellular matrix. Infiltration of tumors by lymphocytes, and in particular CD8+ cytotoxic T cells, is known to predict good prognosis in many types of cancer [1]. T cell infiltration into solid tumors is also associated with increased clinical efficacy of immunotherapies [2]. The beneficial effect of a high abundance of tumor-infiltrating T cells reflects the ability of the immune cells to mount a response against cancer cells [3]. As an important way for the cancer cells to evade immune destruction, tumors can develop a strongly immunosuppressive tumor microenvironment [4–6]. This includes the accumulation of cell types with immunosuppressive activity, such as tumor-associated macrophages (TAMs), myeloid-derived suppressor cells (MDSCs), and regulatory T cells (Tregs) [4]. Upregulation of PD-L1 by cells in the tumor microenvironment constitutes an important and well-studied immune escape mechanism in which the interaction with its receptor PD-1 on T cells lead to inactivation of the T cells [7]. Antibody-mediated blockade of the PD1-PD-L1 interaction has demonstrated remarkable clinical efficacy for many cancer patients and stimulated the research aiming at identifying additional targetable immunosuppressive mechanisms in the tumor microenvironment [7].

In addition to non-malignant stromal cells and immune cells, the tumor microenvironment consists of the extracellular matrix (ECM). Degradation of the ECM surrounding a tumor is an essential part of invasive cancer growth and a main reason for the destruction of the normal tissue [8–10]. Of central importance, the degradation of the ECM is accompanied by the deposition of a different tumor-specific ECM [11]. This new ECM is often of increased density and stiffness, and contains components that are not typically present in the original ECM [12, 13]. Within the last decade, a number of important discoveries have emphasized how the ECM can affect cancer biology [14, 15]. A strong correlation between the density of collagen type I, which is the most abundant component of the tumor ECM, and poor prognosis of breast cancer, gastric cancer and oral cancer has been demonstrated [16–18], and *in vitro* studies have shown that a high-density and stiff ECM can induce a process in epithelial cells resembling malignant transformation [19–21]. Other cell types such as fibroblasts and mesenchymal stem cells have also been demonstrated to respond to the mechanical properties of the surrounding ECM by a process termed cellular mechanosensing [22, 23]. Through the effects on cancer cells and stromal cells, the ECM can augment many of the hallmarks of cancer, such as the induction of angiogenesis [24] and the activation of invasion and metastasis [15, 25]. It remains quite speculative if the ECM can also modulate the immunosuppressive tumor microenvironment and thereby support the cancer’s evasion of immune destruction [15, 26]. It should, however, be noted that the presentation of antigens linked to a stiff surface has been demonstrated to impair TCR-mediated T cell activation, suggesting that T cells possess mechanosensing abilities [27] and others have confirmed that the TCR is affected by mechanical force [28]. Furthermore, tumor-associated remodeling of the ECM, can lead to the deposition of ECM components such as osteopontin, SPARC, versican, and tenascin C, which have been suggested to possess immunosuppressive properties [29–31].

Although cell culture in 3D environments is widely used in the field of cancer biology [32], 3D culture of T cells is less common and has mainly been used for the study of cell migration [33, 34]. The interaction between collagen and tumor-infiltrating T cells was, however, studied by Salmon et al. through the use of elegant *ex vivo* culture of lung tumor slices combined with real-time imaging [35]. Here, collagen fibers were suggested to prevent the migration of T cells from the stromal compartment into the tumor islets. Collagen-density mediated inhibition of directional T cell migration has also been suggested as the reason for intratumoral T cell exclusion in pancreatic cancers [36]. It was not addressed in these studies if the collagen also influences the activity of the T cells. In this study, we employed 3D culture assays to investigate if the collagen-density can directly impact the activity of T cells.

## Materials and Methods

### T cell isolation and culture

Human peripheral blood mononuclear cells (PBMCs) were isolated from healthy donors by gradient centrifugation using Lymphoprep (Alere AS) separation and frozen in fetal calf serum (Sigma Aldrich) with 10% DMSO (Sigma Aldrich). For RNAseq experiments, the PBMCs were enriched for T cells by allowing the cells to adhere overnight and collecting only the non-adherent and loosely adherent cells. For confocal microscopy, T cells were isolated from healthy donors at the Combined Technical Research Core facilities at the National Institute of Dental and Craniofacial Research (NIDCR), NIH using elutriation. For all other experiments, T cells were isolated from PBMCs using magnetic anti-CD3 microbeads (Miltenyi Biotec) according to the manufacturer’s instruction. Cells were cultured at 37°C in a humidified 5% CO_2_ environment in X-vivo media (Sartorius) with 5% human serum (Sigma Aldrich).

### 3D culture in collagen gels

Type I collagen gels were prepared using a modified protocol from Artym and Matsumoto [37]. Briefly, cells were resuspended in a mix of rat tail collagen type I (Corning), 0.02N acetic acid, 10x DMEM with Phenol Red (Sigma Aldrich) and 10x reconstitution buffer (0.2M Hepes (Gibco) and 0.262M NaHCO_3_). To neutralize the pH, 2N NaOH was added to the reconstitution buffer before use. For all cell lines, low density (LD) and high density (HD) gels contained 1 mg/ml and 4 mg/ml collagen type I respectively. First, 300 to 400 μl of the collagen solution was plated per well of a non-tissue culture treated 24-wells plate (Corning) and allowed to polymerize at 37°C for at least 30 min. Afterwards a second layer of collagen solution containing the cells was seeded on top of the first gel and allowed to polymerize for 1-2 h at 37°C, after which 600 μl of culture media was added. For culturing cells in 2D conditions, the same number of cells were seeded in the wells of a tissue culture treated 24-well plate.

### T cell proliferation assay

T cells from healthy donors were transiently stimulated with 10 nM PMA (Sigma Aldrich) and 175 nM ionomycin (Sigma Aldrich) and labelled with CellTrace Violet dye (Thermo Fisher Scientific). 8×10^5^ T cells were seeded in each well within LD or HD collagen gels or on tissue culture plastic. After 5 days of culture in 3D collagen gels or on plastic (2D), cells were treated with 3 mg/ml collagenase (Worthington) solution for 45-60 min at 37C to extract the cells from the gels, washed once with media and once with DPBS and resuspended in FACS buffer containing the following: Live/Dead Fixable Near-IR Dead cell stain (Thermo Fisher Scientific), anti-CD3-APC (cl. SK7), anti-CD4-PE (cl. SK3), and anti-CD8-FITC (cl. HIT8a) (all BD Biosciences). Cells were incubated with the antibodies in the dark at 4°C for 30 minutes, washed twice with FACS buffer, resuspended in FACS buffer and acquired using a BD FACSCanto II flow cytometer (BD Biosciences). Analysis was performed with FlowJo V10 software. Experiments were repeated three times using T cells isolated from different donors.

### Cancer cell proliferation assay

Cell proliferation was determined using the APC BrdU Flow Kit (BD Biosciences) according to the manufacturer’s instructions. Briefly, 50.000 cells were cultured in the different 3D or 2D culture conditions for five days. For labelling, BrdU was added to the medium of each well (final concentration 10 μM) and cells were incubated for 45 min at 37°C in 5% CO_2_. Afterwards, media of the wells with collagen gels was aspirated and replaced with 3 mg/ml collagenase solution (Worthington). Collagenase was also added to the media of 2D cultured cells in concentration similar to which cells in 3D gels were exposed. After complete digestion of the collagen gels, cells were collected and washed once with DPBS (Lonza) and stained with Zombie Aqua Fixable Viability Dye (BioLegend) to determine cell viability. Next, cells were fixed, permeabilized, and stained with APC anti-BrdU antibody according to the manufacturer’s instructions. Cells were resuspended in FACS buffer and acquired using a BD FACSCanto II flow cytometer (BD Biosciences). Analysis was performed with FlowJo V10 software.

### RNA extraction, cDNA synthesis and quantitative real-time-PCR

For RNA isolation, 8×10^5^ PBMCs enriched for T cells or purified T cells (isolated with anti-CD3 microbeads, Miltenyi Biotec) were transiently stimulated with PMA and ionomycin and seeded within LD or HD collagen gels or on tissue culture plastic. After two days, total RNA from cell cultures was purified using RNeasy kit (Qiagen) according to the manufacturer’s instructions. Quality of samples was measured using an Agilent 2100 BioAnalyzer (Agilent Genomics). Afterwards 500 ng – 1 μg RNA per sample was reverse transcribed using iScript cDNA Synthesis Kit (Bio-Rad). The synthesized cDNA was used as template in a real-time quantitative PCR reaction using Brilliant III Ultra-Fast SYBR Green (Agilent Technologies) according to the manufacturer's standard protocol. Equal amounts of cDNA were applied in each reaction mixture. As a control for the specificity of the quantitative real-time PCR, a sample without template was included. The real-time cycler conditions were as follows: initial activation step at 95°C for 3 min, 40 cycles of denaturing at 95°C for 5 s, and annealing/extension at 60°C for 20 s, followed by a melting curve analysis of 65–95°C with 0.5°C increment, 5 s per step.

All measurements were based on triplicates or quadruplicates of each cell culture condition measured in duplicates and normalized to the internal control gene, *ACTB*. Four independent experiments were performed. The comparative cycle threshold (ΔΔCT) method was used to calculate the relative fold changes.

Primers were designed using the Primer-BLAST tool (NCBI, NIH). All primers spanned exon-exon junction, and maximum product length was 250 bp. Primer efficiencies were measured for all primer sets and found to be between 85% and 103%. Primers are listed in Table S2.

### RNAseq

The quantity and purity of isolated RNA were assessed using an Agilent 2100 BioAnalyzer (Agilent Genomics). Total RNA (500 ng) was prepared for sequencing using polydT-mediated cDNA synthesis in accordance with the manufacturer’s (Illumina) instructions. Libraries were made with a NEBNext RNA Library Preparation Kit for Illumina. Library quality was assessed using Fragment Analyzer (AATI), followed by library quantification (Illumina Library Quantification Kit). Sequencing was done on a HiSeq1500 platform (Illumina) with a read length of 50 bp. Sequenced reads were aligned to the human genome assembly hg19 using STAR [38]. Uniquely aligned reads were quantified at exons of annotated genes and normalized to sequence depth and gene length using HOMER [39]. Sequencing depth and alignment information is in Table S3. The number of reads per kilobase per million mapped (RPKM) for all RefSeq annotated genes can be found in Table S1. The analysis of differential expression was performed using DESeq2 package in R [40]. Principal component analysis was performed using R (prcomp package). Heatmaps were generated from z-score normalized RPKM values using R (pheatmap package) on selected sets of genes. MA and Volcano plots were generated using R.

### Cytotoxicity assay

A ^51^Cr-release assay for T cell-mediated cytotoxicity was used to assess the cytotoxicity of tumor infiltrating T cells after 3D culture in collagen matrices of high or low density. Autologous melanoma cells MM33 were used as target cells [41]. Effector cells (T cells) were pre-cultured on plastic or in 3D collagen cultures for 3 days, after which cells were treated with 3 mg/ml collagenase solution for 45-60 min to extract the from the gels, and washed twice with media. Typically, 5×10^5^ target cells in 150 μl RPMI were labelled with 20 μl ^51^Cr (Perkin Elmer) in a 15 ml falcon tube at 37°C for 1 h. After washing, 5×10^3^ target cells per well were plated out in a 96-well plate (Corning) and T cells were added at various effector to target cell (E:T) ratios. Cells were incubated at 37°C for 4 h. Next, the level of ^51^Cr in the supernatant was measured using a Wallac Wizard 1470 automatic γ counter (Perkin Elmer). The maximum ^51^Cr release was determined by addition of 100 μl 10% Triton X-100, and minimum release was determined by addition of 100 μl of RPMI to target cells. Specific lysis was calculated using the following formula: ((cpm _sample_ – cpm _minimum release_)/(cpm _maximum release_ – cpm _minimum release_)) × 100%.

### Statistical analysis

All individual experiments were performed at least three times with at least three replicates per condition and results are presented as mean ± standard error of the mean (SEM) unless otherwise specified. For two-group comparisons between cells from the same donors, paired two-tailed Student’s t-tests were performed. For multi-group comparisons, one-way analysis of variance (ANOVA) was used followed by paired two-tailed Student’s t-tests. All statistical analyses were performed using GraphPad Prism. A p-value <0.05 was considered statistically significant. Correlation between PSR-positive area and CD8+ cell abundance was analyzed by Pearson correlation.

### Additional materials and methods

Detailed information about cancer cell culture, confocal microscopy, flow cytometry analysis of T ell subsets, histology, and ELISA can be found in the Supplementary Data.

## Results

### 3D culture of T cells in different collagen densities impairs proliferation without compromising viability

To investigate if 3D culture in collagen matrices of different collagen concentrations affected the viability of T cells, we isolated T cells from healthy donors and transiently stimulated the cells with PMA and ionomycin. This type of stimulation bypasses T cell receptor activation but acts on several of the same downstream signaling pathways including Protein Kinase C [42]. The T cells were embedded in collagen matrices of high (4 mg/ml) or low (1 mg/ml) collagen concentration, or seeded on regular tissue culture plastic (2D culture) and cultured for 5 days. The selected collagen concentration of 1 mg/ml is representative of healthy normal tissue such as lung or mammary gland whereas 4 mg/ml collagen gels mimic the tissue stiffening occurring in solid tumors [19, 43]. To completely avoid cellular contact with the plastic surface of the wells, the 3D culture was established on top of a pre-generated collagen matrix without cells (Figure 1A). To evaluate if viability of the T cells was affected by the different culture conditions, cells were extracted from the collagen matrices by a brief collagenase-treatment, stained with a live/dead cell marker and analyzed by flow cytometry (Figure 1B). A high viability of more than 95 % was observed in both 2D culture and in 3D culture in different collagen densities. To visualize the 3D culture of T cells in collagen matrices of different collagen concentrations, purified T cells were embedded in collagen matrices and imaged by confocal fluorescent microscopy (Figure 1C-E). As expected, 3D cultured T cells were completely surrounded by collagen (Figure 1C). The structure and density of collagen fibers were clearly different in the matrices of different collagen concentrations but no apparent morphological changes were observed between T cells in 3D culture of high or low collagen density (Figure 1D-E). To examine if T cell proliferation was affected by collagen density, T cells were transiently PMA/ionomycin stimulated, CellTrace Violet (CTV) labeled, and embedded in a high- or low-density collagen matrix or cultured on regular tissue culture plastic. Flow cytometry-based analysis of CTV dilution in CD3+ cells showed a clear reduction in proliferation when cells were cultured in 3D compared to 2D. Interestingly, we also observed a smaller but still significant reduction in proliferation when T cells were cultured in a high-density collagen matrix compared to a low-density collagen matrix (Figure 1F-G). In consistence with other reports [21, 44] we did not observe that proliferation of cancer cell lines was similarly impaired in a high-density collagen matrix (Figure 1H).

**Figure 1.**
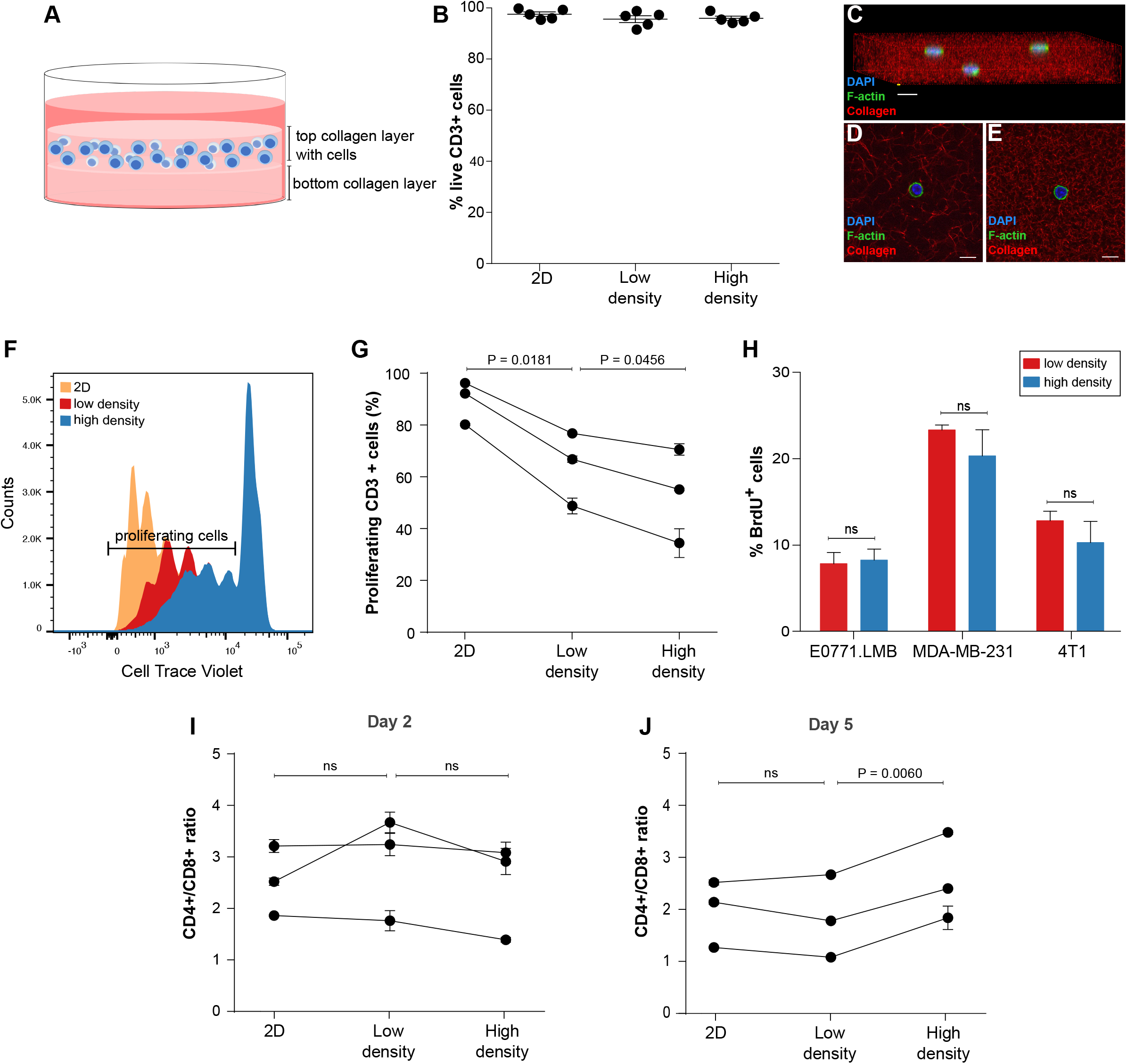
3D culture in high collagen density impairs T cell proliferation. (A) Schematic model of the 3D culture system. (B) T cells were cultured for 5 days in the indicated conditions, and subsequently viability was analyzed by flow cytometry. Each dot in the graph represents an individual donor. Error bars indicate standard error of the mean (SEM). (C-E) T cells cultured in a collagen matrix of low density (1 mg/ml, C-D) or high density (4 mg/ml, E) including fluorescently labeled collagen were imaged by confocal microscopy. (C) 3D projection of collagen matrix with embedded T cells. (D-E) Representative images of individual T cells within a low-density collagen matrix (D) or high-density collagen matrix (E). Size bars: (C-E) 10 μm. (F-G) T cell proliferation after 5 days in culture was measured by flow cytometry based analysis of CellTrace Violet (CTV) dilution. (F) Representative histogram showing CTV dilution in T cells cultured in 2D or in 3D in a low-density collagen matrix or high-density collagen matrix. (G) Quantification of T cell proliferation based on CTV dilution. Three individual donors were analyzed. Connecting lines indicate measurements of the same donor. (H) The breast cancer cell lines EO771.LMB, MDA-MB-231, and 4T1 were cultured in collagen matrices of low or high density for 5 days and analyzed using a BrdU-based flow cytometry assay. The percentage of BrdU-positive cells is depicted. (I-J) The ratio of CD4+ to CD8+ cells was analyzed by flow cytometry after culture for 2 days (I) or 5 days (J). (G-J) Error bars indicate standard deviations of technical replicates.

The effects were similar for CD4+ and CD8+ T cells (Figure S1) and consistently the different culture conditions did not change the ratio of CD4+ cells to CD8+ cells after 2 days in culture (Figure 1I). However, after prolonged 3D culture of T cells for 5 days, the ratio of CD4+ cells to CD8+ cells was higher in a high-density matrix compared to a low-density matrix (Figure 1J), indicating that proliferation and/or survival is slightly more impaired for CD8+ cells than for CD4+ cells in a high-density collagen matrix.

To investigate if the individual effector and memory differentiation subsets of T cells were differentially affected by collagen density, we analyzed the distribution of the different T cell subsets by flow cytometry after 5 days of 3D culture in a high- or low-density collagen matrix or regular 2D culture (Figure S2). 3D culture in a high-density collagen matrix compared to a low-density collagen matrix resulted in a higher fraction of effector memory T cells and a lower fraction of central memory T cells.

To study if the reduced T cell proliferation in a high-density collagen matrix could be reflected in the differential abundance of tumor-infiltrating T cells in breast tumors, we examined 20 samples of resected triple-negative breast cancers, which were all of histological grade 3 and had a diameter between 10 and 20 mm. The samples were immunostained for CD8 and picrosirius red-stained for fibrillar collagen (Figure 2A-D). Consistent with our 3D culture data, samples that contained a high collagen density often had fewer infiltrating CD8+ T cells, although this negative correlation failed to reach statistically significance with this limited material (Figure 2E).

**Figure 2.**
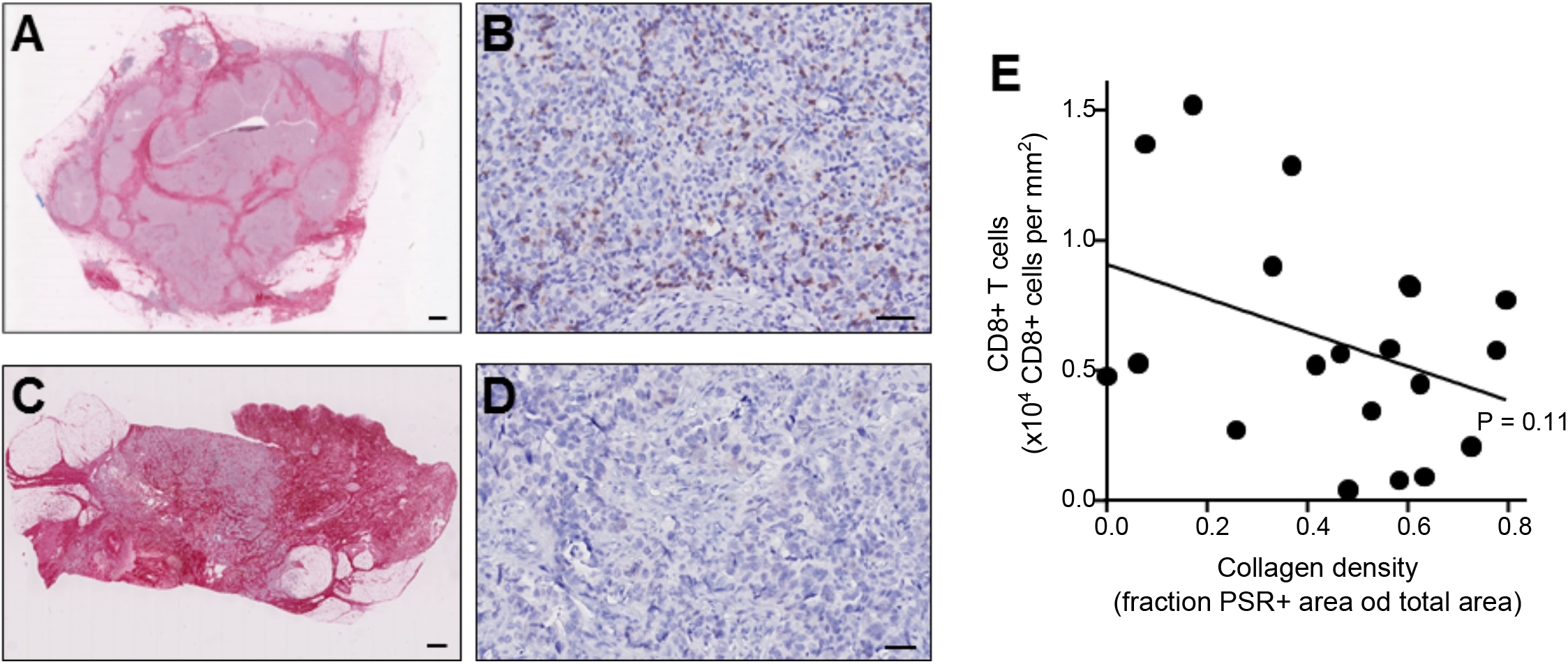
Breast cancer samples of high collagen density have fewer infiltrating T cells. (A-D) Histological sections of triple-negative breast cancers were picrosirius red (PSR)-stained for visualization of fibrillar collagen (A and C, red color) or immunostained for CD8 for visualization of cytotoxic T cells (B and D, brown color). (A-B) Example of a specimen containing low levels of collagen (A) and high abundance of tumor-infiltrating T cells (B). (C-D) Example of a specimen containing high levels of collagen (C) and low abundance of tumor-infiltrating T cells (D). Size bar: (A and C) 1 mm; (B and D) 100 μm. (E) Using Visiopharm-assisted automated image analysis, 20 triple-negative breast cancer sections were analyzed for the PSR-positive area and the number of tumor-infiltrating CD8+ T cells. Each dot in the graph represents an individual cancer sample. Pearson correlation r=0.37, P=0.11.

### The gene expression profile of T cells is regulated by surrounding extracellular matrix

To further elucidate the response of T cells to 3D culture in different collagen densities, T cells were 3D cultured for 2 days in low- or high-density collagen matrices or cultured on regular tissue culture plastic and subjected to RNA sequencing. A principal component analysis shows that the gene expression profile of cells cultured in 3D (low and high density) separate very clearly from the 2D cultured cells (cultured on plastic) and additionally that cells cultured in a high-density collagen matrix cluster separately from cells cultured in a low-density collagen matrix (Figure 3A). The full gene expression dataset is in Table S1. The clear difference between 2D and 3D culture was reflected in 683 differentially regulated genes (FDR < 0.01 and fold change > 1.5) for cells cultured in a low-density collagen matrix compared to regular 2D culture on tissue culture plastic (Figure 3B and Figure S3A) and 1928 differentially regulated genes (FDR < 0.01 and fold change > 1.5) between culture in high-density collagen compared to regular 2D culture (Figure S3B-C). In consistence with the reduced proliferation (Figure 1F-G), downregulated genes were involved in cell cycle processes (Figure 3C and Figure S3E). To investigate if T cell activity was affected by 3D culture, we examined the expression levels of a panel of T cell activity markers (Figure 3D), regulatory T cell (Treg) markers (Figure 3E), and exhaustion markers (Figure 3F). The heatmaps did not show any clear 3D culture-induced changes in T cell activity although some of the genes were indeed significantly regulated, suggesting that 3D culture could influence T cell biology. To identify putative transcription factors (TFs) and TF families that may be responsible for the differential gene expression observed after 3D culture of T cells, we used the computational method ISMARA (Integrated Motif Activity Response Analysis, http://ismara.unibas.ch/fcgi/mara) to model transcription factor activity [45]. Among the top-ranked identified downregulated TF motifs, several known pro-proliferative factors such as Myb and members of the E2F family were included [46, 47] (Figure S4). This is consistent with the gene ontology analysis (Figure 3C) and the observed reduction in cellular proliferation after 3D culture of T cells (Figure 1F-G).

**Figure 3.**
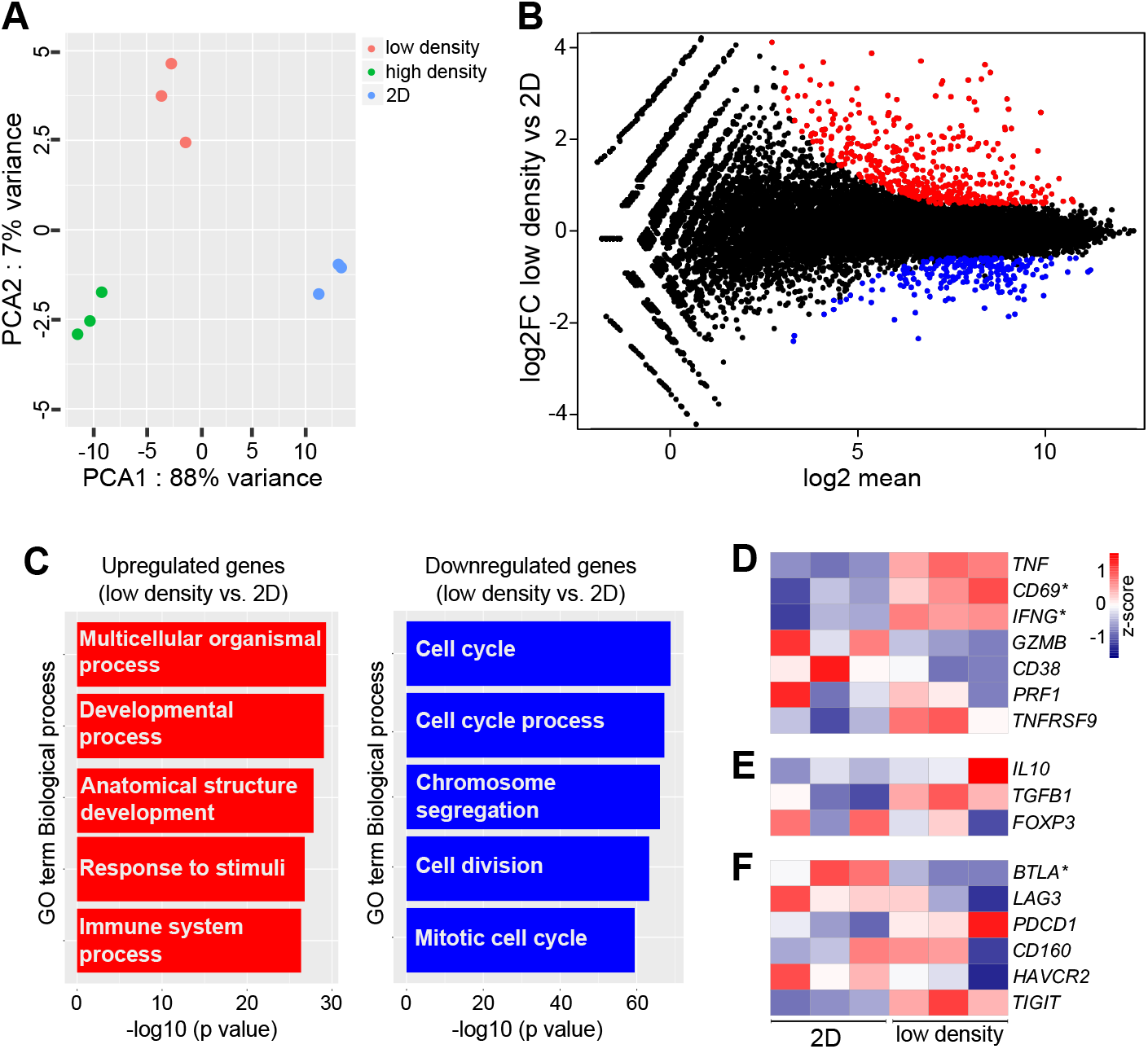
Distinct transcriptomic signatures in 2D culture and in 3D culture in different collagen densities. (A) Principal component analysis of each RNAseq replicate of T cells cultured on plastic (2D) or in 1 mg/ml (low density) or 4 mg/ml (high density) collagen matrices for 2 days. (B) MA plot illustrating the differentially regulated genes (FDR < 0.01 and fold change > +/−1.5) between cells cultured in a low-density collagen matrix or in regular 2D culture. Genes that are upregulated in low-density collagen compared to 2D are shown in red and downregulated genes are shown in blue. (C) Gene ontology analysis illustrates biological processes most significantly enriched within genes that are upregulated (left panel, red bars) or downregulated (right panel, blue bars) in low density collagen compared to 2D. (D-F) Heatmaps of normalized (Z-score) RNAseq read counts of genes encoding markers of T cell activity (D), Tregs (E), or T cell exhaustion (F). (D-F) Asterisks indicate significantly regulated genes.

### T cells cultured in a high-density collagen matrix downregulate markers of cytotoxic activity

To investigate the gene regulation specifically induced by collagen density, we compared T cells cultured in high- vs. low-density collagen matrices and found that 351 genes were differentially expressed (FDR < 0.01 and fold change > 1.5) (Figure 4A and Figure S3D). In alignment with the observed collagen density-induced reduction in proliferation (Figure 1F-G), downregulated genes were involved in cell cycle processes (Figure 4B). Upregulated genes were involved in processes such as chemotaxis and cell migration (Figure 4B). Of the 10 most significantly up- and down-regulated genes (Table 1), several have been suggested to impact T cell activity. CD101, which is upregulated in a high-density matrix, has been suggested to be involved in the negative regulation of T cell activity [48] and the expression levels on Tregs correlate with their immunosuppressive potency [49]. CIP2A, which is downregulated in a high-density matrix, has been suggested to promote T cell activation [50].

**Figure 4.**
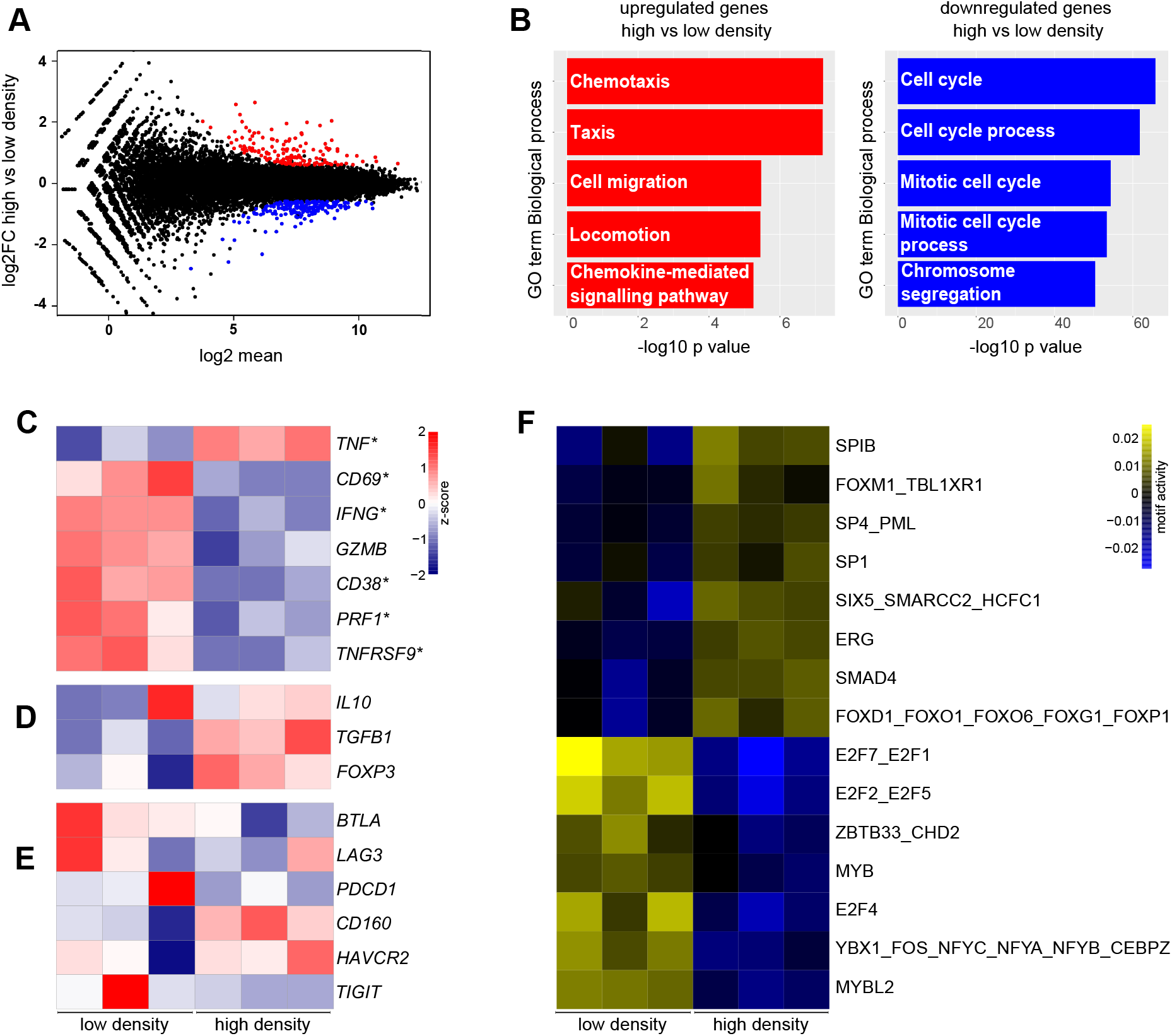
A high-density matrix induces a transcriptomic program indicative of reduced proliferation and cytotoxic activity. (A) MA plot illustrating the differentially regulated genes (FDR < 0.01 and fold change > +/− 1.5) between cells cultured in a high-density collagen matrix or a low-density collagen matrix for 2 days. Genes that are upregulated in high-density collagen compared to low-density are shown in red and downregulated genes are shown in blue. (B) Gene ontology analysis illustrates biological processes most significantly enriched within genes that are upregulated (left panel, red bars) or downregulated (right panel, blue bars) in high density compared to low density. (C-E) Heatmaps of normalized (Z-score) RNAseq read counts of genes encoding markers of T cell activity (C), Tregs (D), or T cell exhaustion (E). (D-E) Asterisks indicate significantly regulated genes. (F) Heatmap of most significantly up- or downregulated TF motifs in high-density vs. low-density collagen.

**Table 1.**
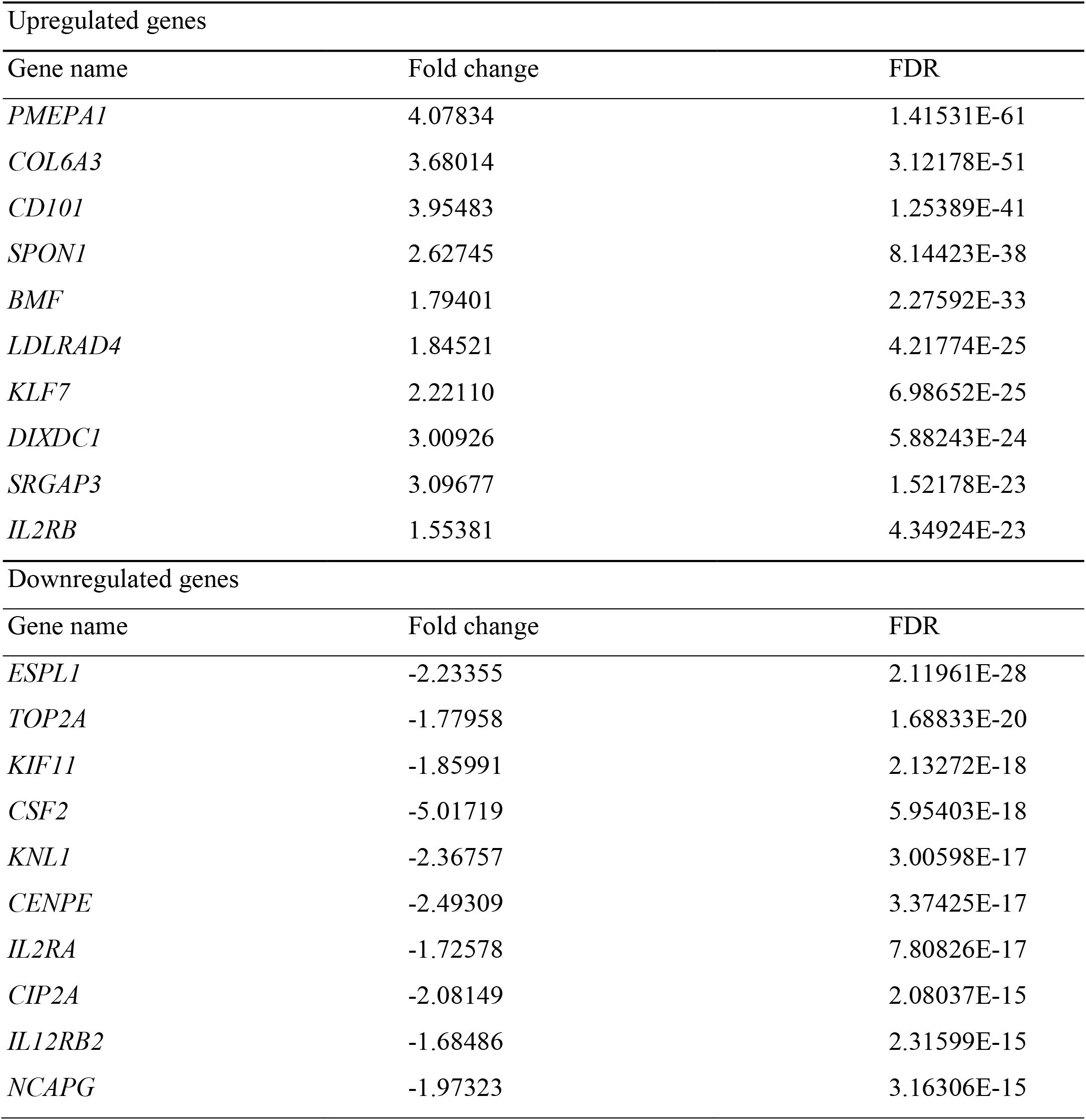
Top up- or downregulated genes in high-density vs. low-density collagen.

| Upregulated genes |  |  |
| --- | --- | --- |
| Gene name | Fold change | FDR |
| *PMEPA1* | 4.07834 | 1.41531E-61 |
| *COL6A3* | 3.68014 | 3.12178E-51 |
| *CD101* | 3.95483 | 1.25389E-41 |
| *SPON1* | 2.62745 | 8.14423E-38 |
| *BMF* | 1.79401 | 2.27592E-33 |
| *LDLRAD4* | 1.84521 | 4.21774E-25 |
| *KLF7* | 2.22110 | 6.98652E-25 |
| *DIXDC1* | 3.00926 | 5.88243E-24 |
| *SRGAP3* | 3.09677 | 1.52178E-23 |
| *IL2RB* | 1.55381 | 4.34924E-23 |
| Downregulated genes |  |  |
| Gene name | Fold change | FDR |
| *ESPL1* | -2.23355 | 2.11961E-28 |
| *TOP2A* | -1.77958 | 1.68833E-20 |
| *KIF11* | -1.85991 | 2.13272E-18 |
| *CSF2* | -5.01719 | 5.95403E-18 |
| *KNL1* | -2.36757 | 3.00598E-17 |
| *CENPE* | -2.49309 | 3.37425E-17 |
| *IL2RA* | -1.72578 | 7.80826E-17 |
| *CIP2A* | -2.08149 | 2.08037E-15 |
| *IL12RB2* | -1.68486 | 2.31599E-15 |
| *NCAPG* | -1.97323 | 3.16306E-15 |

To investigate the potential regulation of T cell activity by collagen density, we visualized the regulation of a panel of cytotoxic activity markers, exhaustion markers, and markers of Tregs (Figure 4C-E). Transcripts encoding six out of seven T cell activity markers were significantly downregulated by culture in high-density collagen compared to low-density collagen (Figure 4C). Only TNF did not follow the same trend and was instead significantly upregulated. At the same time Treg markers were slightly upregulated (Figure 4D). The markers of T cell exhaustion were not systematically regulated by collagen density (Figure 4E). To identify potential transcriptional regulators of these effects, we again used ISMARA to model TF activity. Among the top-ranked downregulated TF motifs, we identified several E2F motifs (Figure 4F), which is consistent with the observed changes in cell proliferation. Among the top-ranked upregulated TF motifs, we identified SMAD4 and FOXO1 motifs (Figure 4F), which are important mediators of TGF-β induced Treg differentiation [51, 52]. In addition, upregulated motifs include PML and FOXM1, which have been suggested to be important for sustaining TGF-β signaling [53, 54].

Altogether, the striking gene expression differences induced by culture in high-density collagen compared to low-density collagen suggests that the T cells acquire a less cytotoxic and more regulatory phenotype.

All of the RNA sequencing experiments were performed using PBMCs isolated from a single healthy donor. To examine if the collagen-density induced gene regulations were donor-specific or reflected a general regulatory mechanism, we examined if similar gene-regulation was observed in cells isolated from four additional healthy donors. A panel of six transcripts, which were significantly regulated by collagen-density in the RNA sequencing experiments (two upregulated and four downregulated), were analyzed by qRT-PCR and compared to the RNA sequencing data (Figure 5A-B). The gene-regulations followed the same pattern as observed in the RNA sequencing experiments, with four of the six genes showing a statistically significant regulation. This experiment as well as the RNA sequencing was performed with PBMCs enriched for T cells by excluding the strongly adherent fraction of the PBMCs, and consequently approximately 70% of the cells were T cells (data not shown). To investigate if the remaining 30% cells, which include myeloid cells, could be critical for mediating this effect of collagen-density we purified T cells from PBMCs from four healthy donors using magnetic anti-CD3 microbeads, resulting in a T cell purity of more than 96% (data not shown). The purified T cells were again transiently PMA-stimulated and cultured in high-density collagen or low-density collagen. Analysis by qRT-PCR of the six-gene panel showed a similar pattern of gene-regulations, indicating that the T cells are directly affected by the surrounding collagen density (Figure 5C).

**Figure 5.**
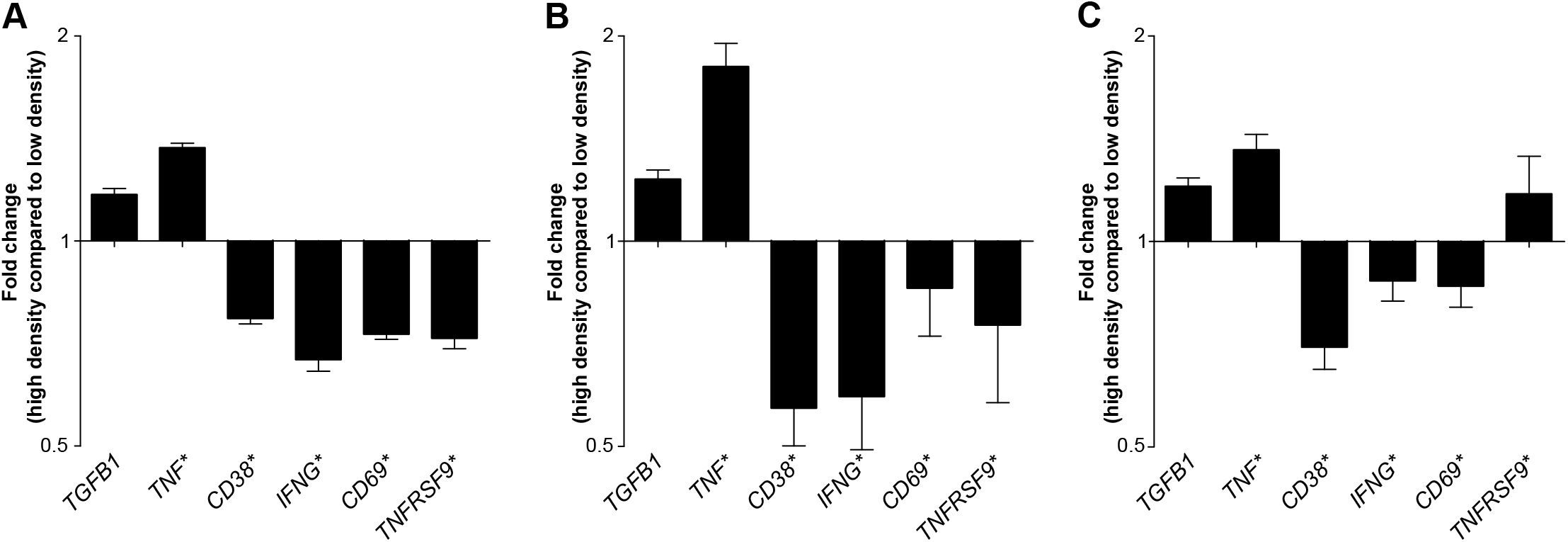
T cells from different donors respond similarly to the surrounding collagen density. (A) Heatmap of normalized (Z-score) RNAseq read counts of a selected panel of differentially regulated genes. (B-C) Heatmap of qRT-PCR analyses of the same panel of genes as in (A) in 4 different donors. Cultured cells were either PBMCs enriched for T cells (B) or purified T cells (C) cultured for 2 days on plastic or in a high-density collagen matrix or a low-density collagen matrix. (A-C) Asterisks indicate significantly regulated genes.

### Collagen density modulates the tumoricidal activity of tumor-infiltrating T cells

To investigate if the collagen density-induced transcriptional regulation of T cells was also reflected in an altered cytotoxic activity, we used a matched set of cultured T cells and melanoma cells isolated from the same tumor fragment. The T cell culture from this patient (MM33) has previously been shown to contain tumor-reactive T cells and to have the ability to lyze autologous melanoma cells [41]. The MM33 T cells were transiently PMA/ionomycin-stimulated and 3D cultured for 2 days in a low-density collagen matrix or a high-density collagen matrix or on regular tissue culture plastic. Subsequently, the cells were extracted from the matrices and assayed for their ability to lyze melanoma cells in a standard four-hour Cr-51 release assay (Figure 6A). The incubation of melanoma cells with increasing numbers of T cells led to increased cell lysis, but the cytotoxic activity of the T cells was impaired after 3D culture compared to 2D culture. Strikingly, cytotoxicity was particularly low for T cells cultured in high-density collagen matrix (Figure 6A-B). The collagen density-induced regulation of T cell cytotoxicity was also reflected in a reduced level of secreted IFN-γ for the MM33 T cells cultured in a high-density collagen matrix (Figure 6C). In these experiments, stimulation with PMA/ionomycin led to noticeable cell death for a large fraction of the cells, which is probably due to the preceding expansion of the cells in high-dose IL-2 containing media. Therefore, we also 3D cultured the T cell for 2 days without any stimulation followed by extraction of cells for cytotoxicity measurements (Figure 6D). In this situation we also observed a reduced cytotoxic activity of the T cells cultured in a high-density collagen matrix compared to the T cells cultured in a low-density collagen matrix (Figure 6D-E). The reduced cytotoxicity was again accompanied by reduced secretion of IFN-γ, although it should be noted that IFN-γ levels were much lower than for the PMA/ionomycin treated T cells (Figure 6F).

**Figure 6.**
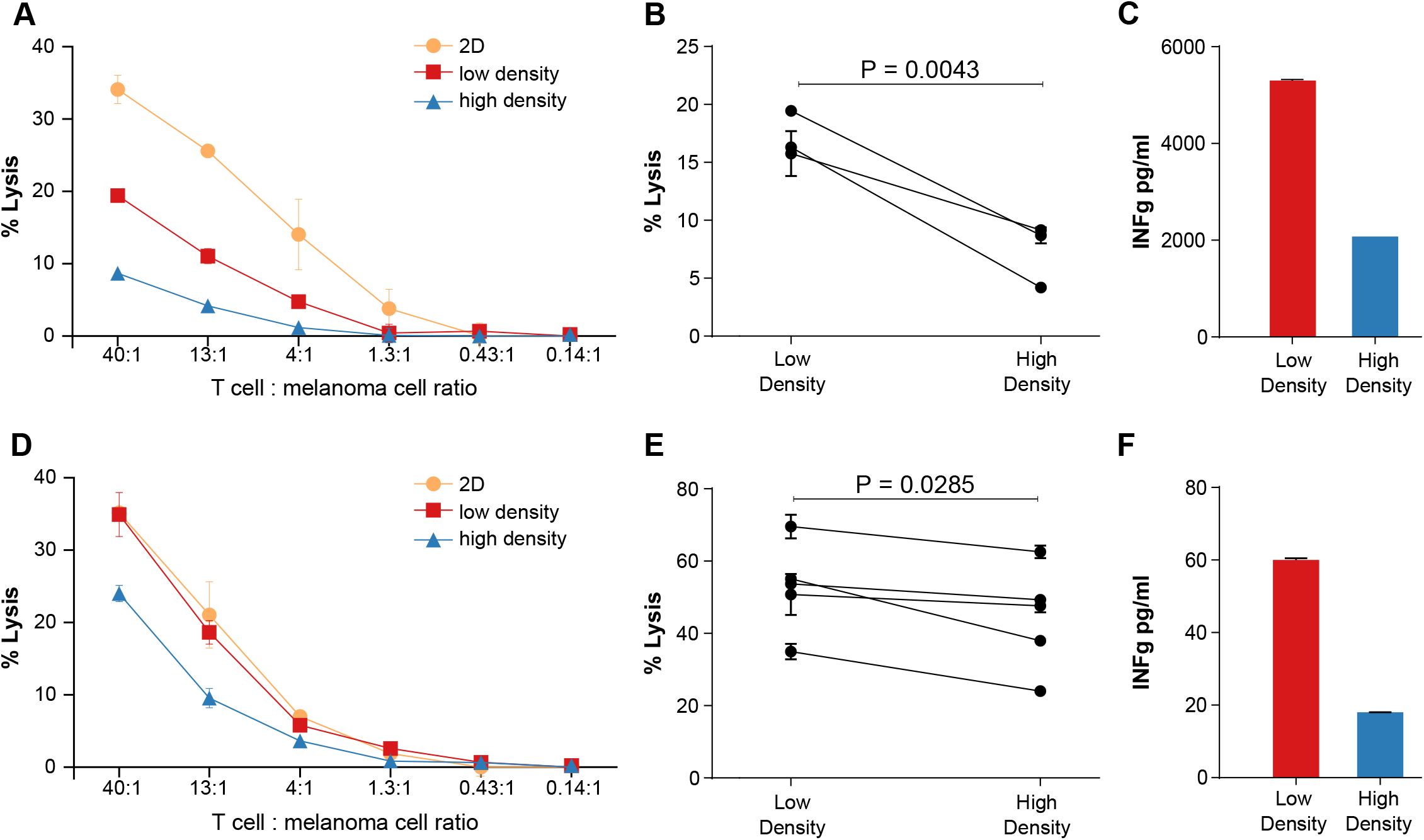
Tumor-infiltrating T cells display reduced cytotoxicity after culture in a high-density collagen matrix. Tumor-infiltrating T cells from melanoma MM33 were cultured for 3 days on plastic (2D) or in collagen matrices of high- or low density, after which the cells were assayed for their ability to lyze autologous MM3 melanoma cells using a 51Cr-release assay. T cells were transiently PMA/ionomycin stimulated before the culture period (A-C) or cultured without any stimulus (D-F). (A and D) Representative example of MM33 melanoma cell lysis after 4 hours incubation with different numbers of T cells, which had been transiently PMA/ionomycin stimulated (A) or directly embedded in collagen (D) and pre-cultured as indicated. (B) Percentage of melanoma cell lysis at the highest T cell: melanoma cell ratio in 3 different experiments. The T cells had been transiently PMA/ionomycin stimulated and cultured for 3 days in a low-density or high-density collagen matrix before extraction and incubation with 51Cr-labeled MM33 melanoma cells. (C and F) IFNγ levels in conditioned media of MM33 T cells, which had been transiently PMA/ionomycin stimulated (C) or directly embedded in collagen (F) and cultured for 3 days in a low-density or high-density collagen matrix. (E) Percentage of melanoma cell lysis at the highest T cell: melanoma cell ratio in 5 different experiments. The T cells had been cultured for 3 days in a low-density or high-density collagen matrix before extraction and incubation with 51Cr-labeled MM33 melanoma cells.

Taken together, our study reveals a novel immunosuppressive mechanism, which could be important for cancer progression and for cancer immunotherapy efficacy.

## Discussion

Collagen density within tumors and their T cell infiltration comprise strong prognostic indicators of poor and good prognosis, respectively. It has, however, not been investigated if these two parameters are independent or if they are interrelated. In this study, we used 3D culture to address whether T cells respond to the surrounding collagen matrix and if the collagen-density alters their activity.

A first important observation made was that T cells indeed respond to their extracellular matrix environment. This was reflected in a dramatically altered transcriptional profile and reduced proliferation for 3D cultured cells compared to regular 2D culture on tissue culture plastic. Gene ontology analysis to identify biological process categories that were statistically enriched confirmed that cell proliferation was significantly affected in 3D. This suggests that 3D culture models, which more accurately mimic tissue environments, could be highly relevant for studies of T cell biology. Although 3D culture of T cells, compared to regular 2D culture, led to a substantial change in the transcriptional profile, inspection of genes encoding markers of activation, Tregs, or exhaustion did not reveal any obvious pattern in the effects on T cell activity.

Another important observation in this study was that T cells clearly respond to the density of the surrounding collagen matrix. Culture of T cells within collagen-matrices of either high or low density resulted in fewer differentially regulated genes compared to the culture of T cells in 2D vs. 3D but nevertheless resulted in striking functional differences. A high collagen-density reduced T cell proliferation and five days of culture in this matrix favored CD4+ T cells over CD8+ T cells. A similar reduction in cell proliferation was not observed for three different breast cancer cell lines, suggesting that within a tumor of high collagen density, T cell proliferation (especially CD8+ T cell proliferation) is impaired whereas cancer cells are unaffected. In line with these observations, we observed in a limited number of triple-negative breast cancer samples, that collagen density seemed to negatively impact CD8+ T cell abundance. The effect of collagen density on T cell proliferation could constitute a new immunosuppressive mechanism within the tumor microenvironment and provide an explanation for the correlation between collagen density in tumors and cancer patient prognosis.

In alignment with the effect of the surrounding collagen density on T cell proliferation, inspection of differentially regulated genes confirmed that cell cycle processes were affected. In addition, markers of cytotoxic T cell activity were clearly downregulated by a high-density collagen matrix compared to a low-density matrix whereas markers of Tregs were upregulated. This striking observation suggests that collagen density, in addition to reducing T cell proliferation, impairs cytotoxic activity. Analysis of putative transcription factor motifs, which could mediate the observed transcriptional changes, identify decreased activity of E2F motifs and increased activity of transcription factor motifs downstream of TGF-β signaling. These findings suggest that E2F transcription factors could be involved in the collagen density-induced inhibition of T cell proliferation and that autocrine TGF-β signaling could be centrally engaged in the modulation of T cell activity.

Using a unique T cell culture and melanoma cell culture established from the same resected melanoma, we could also demonstrate that the tumor-infiltrating T cells were indeed less efficient at killing the melanoma cells after culture in a high-density collagen matrix compared to a low-density collagen matrix. This observation suggests that the collagen-density of the tumor microenvironment can support the cancer cells’ escape from immune destruction by reducing T cell proliferation and by modulating the cytotoxic activity of tumor-infiltrating T cells. This identified immunosuppressive mechanism could be of relevance during tumor progression but also have importance for the efficacy of cancer immunotherapy.

Although this is the first study to directly assess the response of T cells to the surrounding collagen density, the potential of collagen to modulate immune activity is supported by a study of tissue regeneration, in which collagen implantation in wounded muscles of mice was shown to induce an immunosuppressive microenvironment [55]. This effect involved M2-polarization of macrophages, which led to Th2-polarization of T cells. In our study we focused on the ability of the surrounding collagen to directly regulate T cell activity, but it is possible that collagen-induced M2-polarization of macrophages could further augment this modulation of T cells.

In addition to the effects of collagen-density on T cell activity observed in this study, others have suggested that stromal collagen could limit the migration of T cells into the tumor islets and thereby impede their contact with cancer cells [35, 36]. The impaired T cell migration into tumor islets was suggested to be caused by reduced motility in collagen-dense region combined with inappropriate guidance of T cells by the collagen fibers aligned in parallel to tumor islets. These studies further underscore that collagen within the tumor microenvironment could be an important regulator of anticancer immunity.

## Conclusion

By using 3D culture of T cells, we have identified collagen density as a novel regulator of anti-cancer T cell activity. This immunosuppressive mechanism could be of central importance for the cancer cells’ evasion of immune-destruction and could constitute a novel therapeutic target for enhancing immunotherapy efficacy.

## Supporting information

## Acknowledgments

We thank Drs. Thomas H. Bugge and Marie Kveiborg for critically reviewing this manuscript.

## Author contributions

Conceptualization, D.H.M. and D.E.K.; Methodology, D.H.M., D.E.K., L.H.E, M.D., I.M.S., P.t.S. and L.G.; Investigation, D.H.M., D.E.K., A.M.H.L., M.C., A.K., M.S.S., A.M.C.S., A.R., L.G.; Writing – Original Draft, D.H.M. and D.E.K.; Writing – Review & Editing, D.H.M., D.E.K., A.M.H.L., M.C., A.K., M.S.S., A.M.C.S., A.R., L.H.E., M.D., I.M.S., P.t.S., L.G.

## Availability of data and material

The datasets obtained in the current study are available from the corresponding author or available from the indicated sources.

## Consent for publication

Not applicable

## Ethics approval and consent to participate

Healthy donor PBLs were obtained from buffy coats available from the central blood bank of the capital region of Copenhagen and informed consent was obtained from all donors. Breast cancer samples were anonymized and used upon approval by the Scientific Ethics Committee for The Capital Region of Denmark.

**Abbreviations**
ECM: Extracellular matrix
MDSCs: Myeloid-derived suppressor cells
PBMCs: Peripheral blood mononuclear cells
PMA: Phorbol 12-myristate 13-acetate
TAMs: Tumor-associated macrophages
PD-1: Programmed cell death protein 1
PD-L1: Programmed death-ligand 1
TGF-β: Transforming growth factor β
CTV: Cell trace violet
TCR: T cell receptor
Tregs: regulatory T cells

