## Supplementary material for "Collagen density regulates the activity of tumor-infiltrating T cells"

**Supplementary Materials and Methods:**

*Cancer cell culture*

All cancer cells were cultured in RPMI-1640 (Life technologies) with 10% fetal calf serum and 1% Penicillin/Streptomycin (Thermo Fisher Scientific) and cell cultures were split 2 to 3 times a week. Cells were cultured in a humidified 5% $CO_2$ incubator at 37ºC. Cancer cell lines 4T1 and MDA-MB-231 were obtained from American Type Culture Collection (ATCC) and the EO771.LMB breast cancer cell line was a kind gift from Dr. Robin L. Anderson, University of Melbourne. MM33 melanoma cells were isolated from a resected tumor, which had also been used for establishment of a T cell culture as previously described (1). Cell lines were tested negative for mycoplasma using the PCR Mycoplasma Test Kit (AppliChem) less than a year before the use for experiments. Cells were used for experiments within 10 passages after thawing.

*Confocal Microscopy*

T cells were rested overnight in X-vivo + 5% human serum and embedded in three-dimensional collagen gels of 1 and 4 mg/ml collagen prepared as described above (see 3D culture in collagen gels), except additionally 10% Alexa flour 647-labelled collagen type I was incorporated into each gel. The collagen type I with Alexa flour 647 was done as previously described (2). 200 μl of the gel mixed with a total of $8x10^5$ T cells was pipetted into each well of a glass bottom 24-well, No. 0 Coverslip, 13 mm Glass Diameter (MatTek Corporation) and allowed to polymerize at 37ºC, 5% $CO_2$ for 45 min. Thereafter, 500 μl of X- vivo + 5% Human Serum was added on top of the gels and the cells were cultured overnight. Prior to microscopy, samples were stained as described (3). Briefly, the samples were fixed with 4% formaldehyde, 5% sucrose for 1 h and permeabilized with 0.5% Triton-X for 10 min. Next, F-actin was stained for 1 h with 1:40 diluted Alexa Fluor 488 Phalloidin (Thermo Fisher Scientific) in PBS, and the nuclei were stained for 20 min with 1:1000 diluted DAPI (Sigma Aldrich) in PBS. Confocal imaging was performed using an IX83 confocal microscope (Olympus) equipped with a FluoView 1200 scanning head (Olympus). All images were acquired using an Olympus UPlanSApo 60x Oil Objective. Images were processed using ImageJ (NIH).

*Flow Cytometry analysis of T cell subsets*

Purified T cells from PBMCs isolated from three healthy donors were cultured on plastic (2D) or in collagen matrices (3D) of low or high density for five days. T cells from individual donors were extracted from the collagen gels and analysed by flow cytometry. Between 2–6 × $10^5$ cells were stained with a mix of the following fluorochrome-conjugated monoclonal antibodies: anti-CD3-BV510 (cl. SK7), anti-CD4-PerCP (cl. SK3), anti-CD8-BV421 (cl. RPA-T8), anti-CD45RA-FITC (cl. L48), and anti-CD62L-APC (cl. DREG-56) (all BD Biosciences). For dead cell exclusion, Live/Dead Fixable Near-IR Dead cell stain - NIR (Thermo Fisher Scientific) was included. Staining was performed for 20–30 minutes protected from light, at 4°C. Cells were washed once prior to and twice post staining with DPBS with 0.1% bovine serum albumin and 5mM ethylenediaminetetraacetic acid (FACS Buffer). Cells were resuspended in FACS buffer and acquired using a BD FACSCanto II flow cytometer (BD Biosciences). Analysis was performed with FlowJo V10 software. Experiments were repeated three times using T cells isolated from different donors.

*Histology*

Formalin-fixed paraffin-embedded tissue specimens from 20 patients with triple-negative ductal mammary carcinoma were included in this study. Tumors were histological grade 3, between 10 and 20 mm in diameter, and were resected from patients with non-metastatic disease. For immunostaining and picrosirius red-staining, sections were deparaffinized with xylene and hydrated through ethanol/water dilutions, and subjected to a pretreatment with TEG (10mM Tris, 0,5mM EGTA, pH 9,0) for 15 minutes at 98°C. Immunohistochemistry using mouse monoclonal anti-CD8 antibody (1:100 dilution, DAKO) was performed by incubating sections with primary antibodies overnight at 4°C and for 45 minutes at room temperature with EnVision+ System Labeled Polymer–horseradish peroxidase (HRP) anti-mouse (DAKO), according to manufacturer's instructions. Chromogen staining was performed using NovaRED HRP substrate kit (VWR International). All antibodies were diluted in Antibody Diluent with Background Reducing Components (DAKO). Counterstaining was performed using Harris haematoxylin (Histolab). For detection of fibrillar collagen, sections were stained with 0.1% Sirius red diluted in saturated picric acid (Ampliqon) and counterstained with Weigert’s hematoxilin.

Stained sections were scanned using a Panoramic MIDI slide scanner (3D Histech). Using Visiomorph software, the picrosirius red-positive area of each specimen was determined. For the automated detection of the number of CD8-positive cells, region of interests comprising the tumor tissue (excluding surrounding stroma or fat) were marked based on H&E staining of adjacent sections. The number of CD8-positive cells were detected using Visiopharm software.

*ELISA*

Cell culture supernatants were analyzed using IFN gamma Human Uncoated ELISA Kit (Thermo Fisher Scientific) according to the manufacturer's instructions. Optical density or absorption at 450 nm was measured using the Epoch microplate spectrophotometer (BioTek) and analyzed using ELISA Gen5 software (BioTek). Conditioned media was collected from quadruplicates of each condition and plated in duplicates and the mean absorbance was calculated from the standard curve.
