## Supplementary material for "Collagen density regulates the activity of tumor-infiltrating T cells"

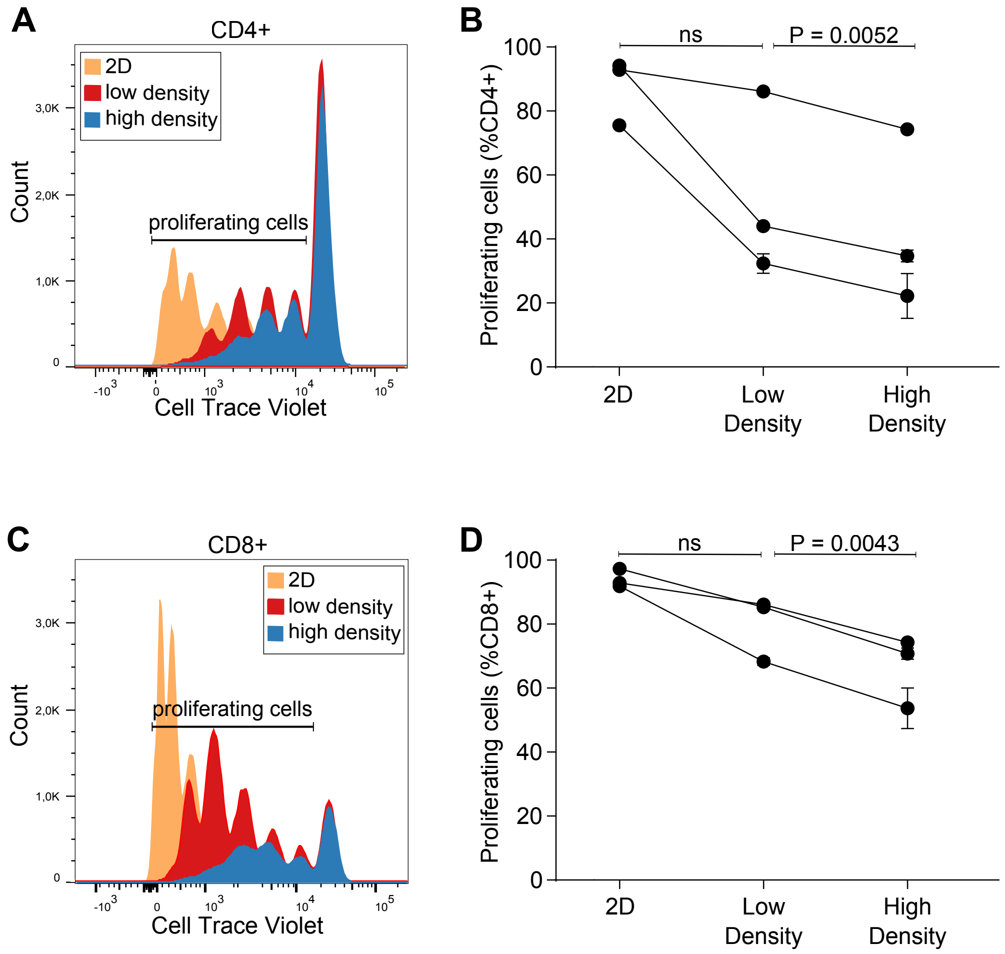


Figure S1. Proliferation of CD4+ and CD8+ T cells
Proliferation of CD4+ and CD8+ T cells after 5 days in culture was measured by flow cytometry based analysis of CellTrace Violet (CTV) dilution. (A and C) Representative histogram showing CTV dilution in CD4+ T cells (A) or CD8+ T cells (C) cultured in 2D or in 3D in a low-density collagen matrix or high-density collagen matrix. (B and D) Quantification of CD4+ T cell (B) or CD8+ T cell (D) proliferation based on CTV dilution. Three individual donors were analyzed. Connecting lines indicate measurements of the same donor.

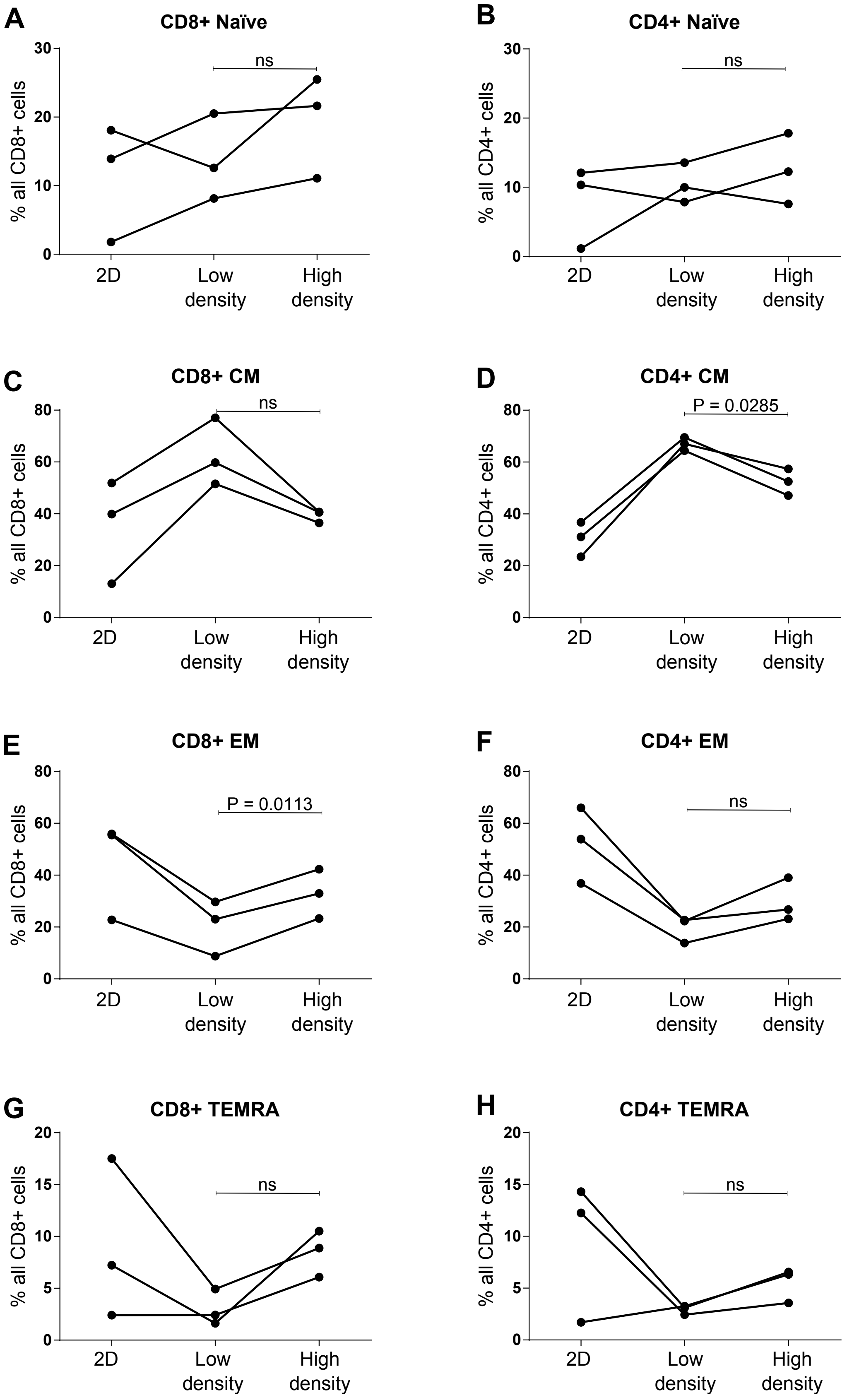


Figure S2. Fraction of T cell subsets after 2D culture or 3D culture in different collagen densities.

T cells were cultured for 5 days on plastic (2D) or in collagen matrices of high- or low density and analyzed by flow cytometry. CD8+ cells (A, C, E, G) and CD4+ cells (B, D, F, G) were divided into Naïve (CD45RA+,CD62L+ (A-B)), Central Memory (CD45RA-,CD62L+ (C -D)), Effector Memory (CD45RA-,CD62L- E-F)), or TEMRA cells (CD45RA+,CD62L- (G-H)). Three individual donors were analyzed. Connecting lines indicate measurements of the same donor.

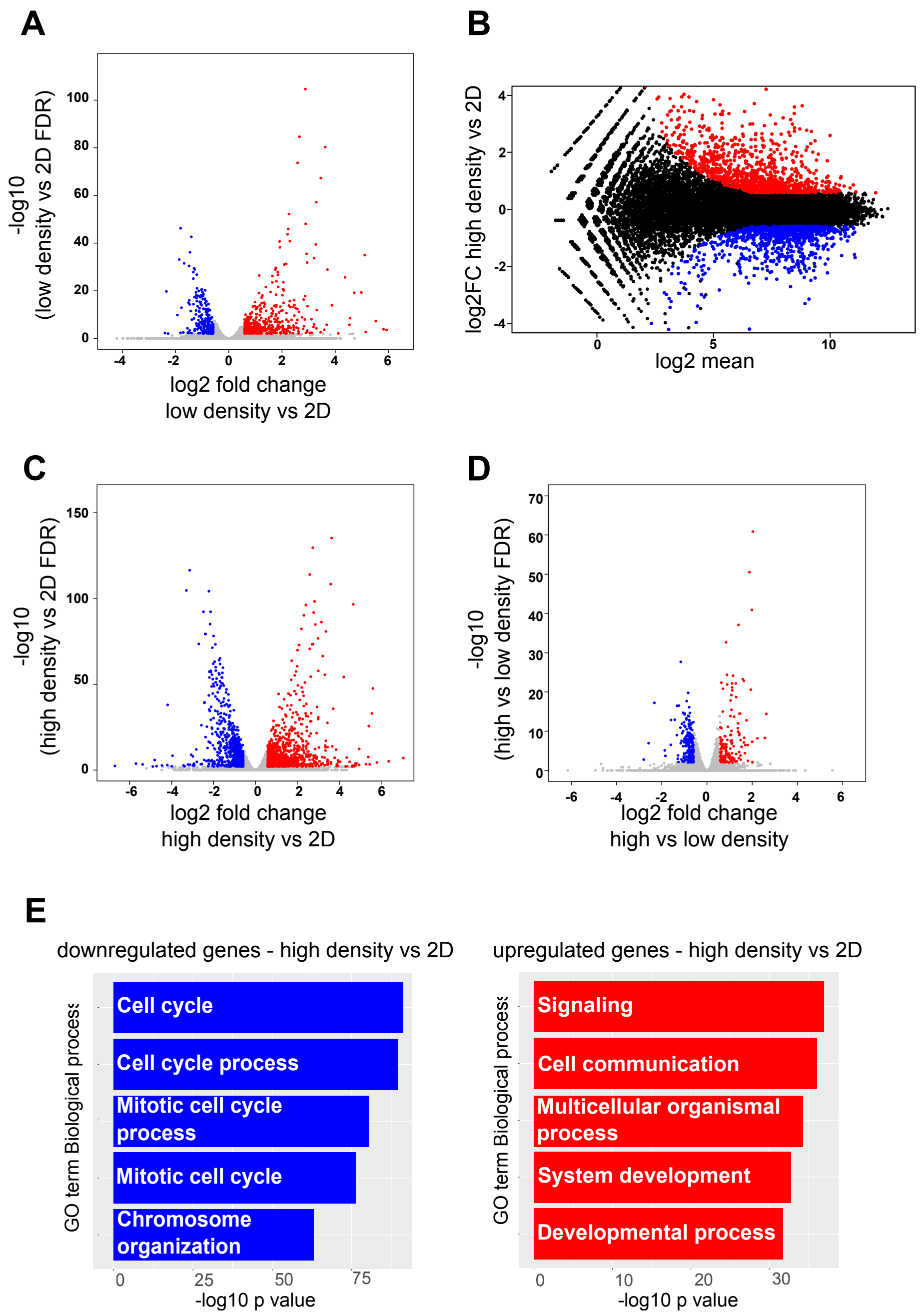


Figure S3. RNAseq data

(A, C-D) Volcano plots illustrating the differentially regulated genes (FDR < 0.01 and fold change > +/- 1.5) between cells cultured in a low-density collagen matrix compared to regular 2D culture (A), high-density collagen matrix compared to regular 2D culture (C), or high-density collagen matrix compared to low-density collagen matrix (D). (B) MA plot illustrating the differentially regulated genes (FDR < 0.01 and fold change > +/- 1.5) between cells cultured in a low-density collagen matrix or in regular 2D culture. (A-D) Genes that are upregulated in are shown in red and downregulated genes are shown in blue. (E) Gene ontology analysis illustrates biological processes most significantly enriched within genes that are upregulated (left panel, red bars) or downregulated (right panel, blue bars) in high-density collagen compared to 2D.

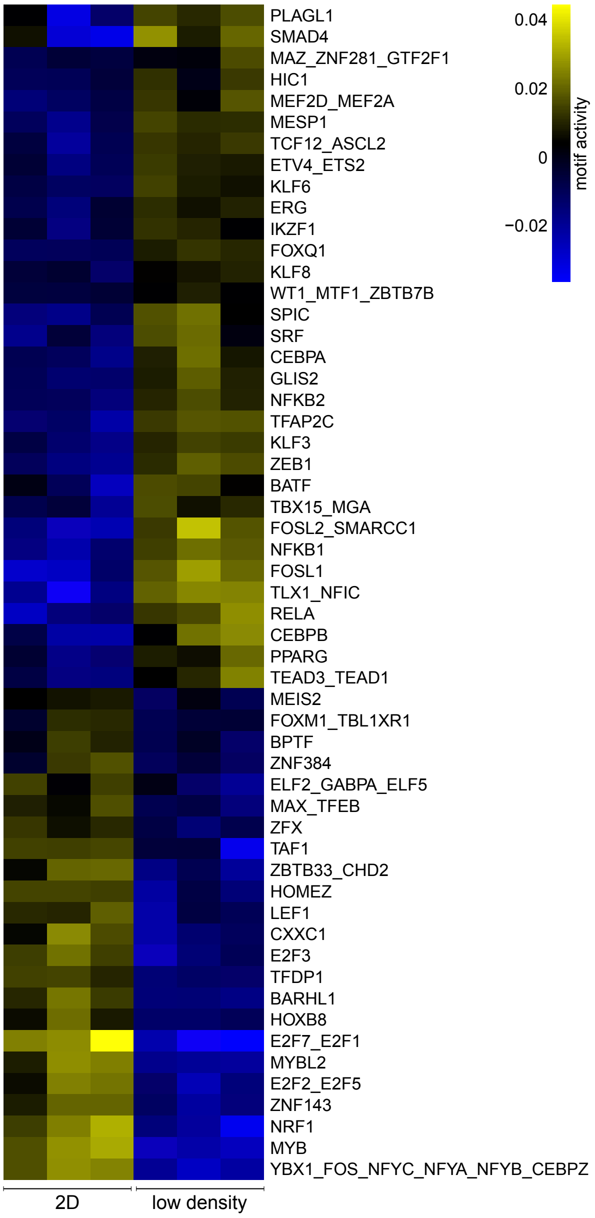


Figure S4. Regulated transcription factors (TF) motifs after 2D culture or 3D culture in low density collagen.

Heatmap illustrating the most significantly up- and downregulated TF motifs in low-density collagen vs. regular 2D culture based on ISMARA analysis. Motifs with Z-values > 1.5 were included in the heatmap

Table S2. Sequences of primers used for RT-qPCR. All primers were designed using the NCBI gene database and the primer-BLAST tool.

| Gene | Protein | Primer direction | Sequence (5' to 3’) |
|---|---|---|---|
| *ACTB* | β-actin | Forward | AGAGCTACGAGCTGCCTGAC |
| | | Reverse | AGCACTGTGTTGGCGTACAG |
| *CD69* | CD69 | Forward | GATGCCACCAGTCCCCATTT |
| | | Reverse | TGGCCCACTGATAAGGCAATG |
| *CD38* | CD38 | Forward | GGTTTCCCGCAGGTTTGC |
| | | Reverse | TCCACACTCCCAAAAGTGCT |
| *TNFRSF9* | C137 | Forward | TGCAGGCAGTGTAAAGGTGT |
| | | Reverse | TCGACAGATGCCACGTTTCT |
| *INFG* | INFγ | Forward | TCGTTTTGGGTTCTCTTGGCT |
| | | Reverse | TTTTCTGTCACTCTCCTCTTTCCA |
| *TNF* | TNFα | Forward | AGCCCATGTTGTAGCAAACCC |
| | | Reverse | GGACCTGGGAGTAGATGAGGT |
| *TGFB1* | TGFβ | Forward | CTGGCGATACCTCAGCAACC |
| | | Reverse | GTGAACCCGTTGATGTCCACTT |

Table S3. RNA sequencing depth and alignment info

| Sample | organism | ref genome | Number of input reads | Uniquely mapped reads | Uniquely mapped reads % |
|---|---|---|---|---|---|
| High-density rep1 | human | hg19 | 16316259 | 12987692 | 79,60% |
| High-density rep2 | human | hg19 | 20827452 | 16565647 | 79,54% |
| High-density rep3 | human | hg19 | 15690290 | 12488770 | 79,60% |
| Low-density rep1 | human | hg19 | 17806762 | 14191720 | 79,70% |
| Low-density rep2 | human | hg19 | 14490086 | 11605314 | 80,09% |
| Low-density rep3 | human | hg19 | 11194136 | 8966651 | 80,10% |
| 2D rep1 | human | hg19 | 14217463 | 11462456 | 80,62% |
| 2D rep2 | human | hg19 | 11640177 | 9254300 | 79,50% |
| 2D rep3 | human | hg19 | 12375402 | 9929856 | 80,24% |
